# Glycan Painting: Triplex Lectin Staining Enables Visualization of Cell-Type-Specific Glycan Profiles in Tissue Sections

**DOI:** 10.64898/2026.04.27.720979

**Authors:** Akira Nagasaki

## Abstract

Multiplex staining is a technique that allows the identification of cell types within a single tissue section by simultaneously detecting multiple molecular markers. Generally, multiplex staining is performed using several combinations of probes, including specific antibodies, nucleic acid probes, and lectins. Here, a novel multiplex staining strategy that relies exclusively on lectin probes that target glycans is presented. Glycans have a vast variety of structural forms that vary depending on cell type-specific modifications. Furthermore, an enormous number of glycan-binding molecules, collectively known as lectins, exist in the biological world. Each lectin displays specificity for a particular glycan motif while maintaining broad affinity. Although lectin-based cell staining has been used in various applications, the partial and limited specificity of lectins has hindered the use of glycan-targeted multiplex staining with lectins. In addition, lectin probes have largely been avoided for cell-type identification because of the absence of strict cell-type-specific glycans. Here, a novel staining method, Glycan Painting, is introduced. Rather than viewing the partial specificity of lectins and the broad, non-cell-type-specific distribution of glycans as drawbacks, this approach turns these features into advantages by generating distinct color patterns that comprehensively visualize cell-type-specific glycan combinations and enable full-color imaging of tissues.

## Introduction

Biological tissues are composed of multiple cell types and states arranged in complex spatial patterns. A comprehensive understanding of the spatial arrangement of all cell types within an organism is essential for elucidating the nature of life. Multicolor staining based on immunostaining can distinguish different cell populations and visualize their spatial relationships. While antibody stripping methods recently allowed the sequential detection of dozens of different antigen molecules on single tissue sections, as demonstrated in cyclic immunofluorescence approaches (Semba and Ishimoto 2024), conventional fluorescence microscopy is generally limited to the detection of approximately three or four spectrally separable fluorescence channels. However, there are some unavoidable challenges when performing multiple antibody staining, including antibody quality and reproducibility, antigen retrieval conditions, and overlapping excitation and emission spectra of fluorescent dyes (Harms et al. 2023).

Glycans collectively refer to carbohydrate chains composed of multiple monosaccharides linked together by glycosidic bonds and are widely present in both eukaryotes, including animal and plant cells, and prokaryotes. It is known that glycans are complex carbohydrate structures that bind to proteins and lipids, play important and essential roles in cellular identify, mediating cell-cell interaction, regulating development, and protein functions (He et al. 2024). On the other hand, a vast number of glycan-binding molecules, collectively known as lectin, exists in biological systems (Manning et al. 2017). To analyze the functions of glycans, fluorescent dye-labeled lectins have been widely used as probes for the detection of glycans. Fluorescent lectins enable visualization of glycan distribution at the cellular and tissue levels and the analysis of cell-type-specific glycosylation patterns in histofluorescent and flow cytometric applications (Laughlin and Bertozzi 2009). However, the limited and overlapping specificity of lectins for glycans is considered as a major challenge (Mattox and Bailey-Kellogg 2021; Bojar et al. 2022; Brooks 2024). Moreover, despite the vast diversity of glycans, none exhibit strict cell type specificity (Tao et al. 2008), which has historically made the development of glycan-targeting cell markers difficult. These limitations have hindered the development of multiplex lectin-based staining methods.

Here, focus is placed on the unique combinations and relative proportions of cell-type-specific glycans to define each cell type within tissues of living organisms. Merged images by triplex fluorescent staining of tissue sections using three different types of lectins with distinct fluorescent dyes enabled the assignment of unique colors to individual cell types in tissues based on the combination of three probes. Mouse embryo sections stained with Lectin Painting revealed unique full-color images that have never been previously achieved using conventional staining methods. Traditionally, lectin staining provides less information than antibody-based approaches for the identification of cell type. However, Glycan Painting demonstrates that even simple lectin-based approaches can provide sufficient information to overcome the limitations of conventional lectin staining.

## Materials and Methods

### Glycan Painting

Cells were cultured on collagen-coated coverslips that had been coated with a 0.001% (w/v) solution of type I collagen (Research Institute for Functional Peptides, Yamagata, Japan). Cells were fixed with 4% paraformaldehyde in phosphate-buffered saline (PBS) for 20 min and permeabilized with ethanol for 5 min. After washing with PBS containing 0.1 g/mL CaCl_2_ and 0.1 g/mL MgCl_2_, hereafter referred as PBS (+), cells were stained with concanavalin A (ConA) CF405 (Fujifilm, Tokyo, Japan), wheat germ agglutinin (WGA) CFR488 (Fujifilm) and *Lycopersicon esculentum* (tomato) lectin Dylight 594 (Thermo Fisher Scientific) in PBS (+) for 60 min at room temperature. After washing, the cells were fixed again with 10% formalin natural buffer solution (Fujifilm) for 10 min and mounted in a glycerol based-mounting medium containing 1.25% w/v 1,4-diazobicyclo{2.2.2}octane (DABCO, Fujifilm).

Formalin-fixed, paraffin-embedded mouse embryo tissue sections were purchased from Genostaff (Tokyo, Japan). Deparaffinization was performed with PathoClean (Fujifilm) and ethanol. The deparaffinized sections were washed 1xTBS (Fujifilm) containing 0.1 g/mL CaCl_2_and 0.1 g/mL MgCl_2_(TBS (+)) and blocked with TBS (+) containing 1x carbo-free blocking solution (Vector Laboratories, CA, USA). After washing with TBS (+), sections were stained with ConA CF405 (Fujifilm), WGA CFR488 (Fujifilm) and LEL Dylight 594 (Thermo Fisher Scientific) in TBS (+) for 90 min at room temperature. Stained samples were washed with TBS (+) three times and refixed with 10% formalin natural buffer solution (Fujifilm) for 10 min. Finally, stained tissue sections were mounted with polyvinyl alcohol based-mounting medium (75% (v/v) diluted laundry glue liquid, Kaneyo Nol (Kaneyo Soap, Tokyo, Japan) containing 15 mg/mL DABCO, 50 mM Tris-HCl (pH 7.5), and 20% (w/v) glycerol (Fujifilm) (final concentrations). Fluorescence images were acquired using an IX83 fluorescence microscope (Evident, Tokyo, Japan) and a Nikon AX confocal microscope (Nikon, Tokyo, Japan). Acquisition conditions for each image are described in the corresponding figure legends.

Hematoxylin and eosin (H&E) staining of the mouse embryo tissue sections was performed using an H&E staining kit according to the manufacturer’s instructions (ScyTek Laboratories, UT, USA).

## Results and Discussion

Multicellular organisms are comprised of several hundred distinct cell types classified according to their morphology and function. H&E staining is a widely used and standard method for visualizing colorless and transparent cells in tissue sections (Morrison et al. 2022). In this approach, cell identification relies on morphological features such as nuclear shape, cell size, and tissue architecture. Therefore, this method makes it difficult to distinguish cell types in complex tissues and among morphologically similar cell populations. If a technology could distinguish specific cell populations based on color, it would represent a major advance in fundamental biology research and the field of pathology. In this study, focus was placed on the distinct combinations of glycans present in different cell types. Glycans, which are covalently bound to proteins and lipids, play an important role in various cellular events, including cell recognition, immune response and cell adhesion. Glycan structures are generated through enzymatic biosynthesis, involving enzymes such as glycosidases and glycosyltransferases, resulting in remarkable structural diversity and substantial variation in glycan composition among different cell types. Although a large number of glycan types exist in living organisms, identifying strictly cell type-specific glycans remains difficult. In contrast, the combinations of glycans present in cells are unique to each cell type. Based on this concept, it was hypothesized that extracting information on cellular glycan profiles, which reflect the unique glycan combination of each cell type, would enable the discrimination of all cells within tissue sections. Cellular glycan profiles were obtained by merging fluorescence images acquired from triplex staining using three lectins, each labeled with a different fluorescent dye. The merged images revealed cell-type-specific glycan combinations as distinct colors corresponding to their unique glycan combinations. This method is based on the three primary colors of light and generates full-color images through the additive combination of red, green, and blue fluorescence, analogous to the mechanism of an LCD monitor (Supplementary Figure 1). Consequently, each cell type acquires a specific color signature. Furthermore, given that mammals are estimated to consist of a few hundred cell types, an 8-bit color palette would theoretically allow distinct color assignment to a large population of cell types.

First, the triple staining of fluorescent lectin at the cellular level was examined. Figure 1A shows the images of several cell lines stained with three types of fluorescent dye-labeled lectins: ConA (Sumner and Howell 1936), WGA (Nagata and Burger 1974) and LEL (Nachbar et al. 1980), which conventionally recognize mannose, GlcNAc and poly-GlcNAc, respectively. These stained images represent composite colors generated from three fluorescent lectins, with each cell type displaying a distinct color, indicating a cell-type-specific combination of glycans. The amount of glycans varies greatly between cell types, so it is difficult to adjust the contrast and intensity of each image on the same screen. Therefore, auto-contrast was performed for each image individually using FIJI ImageJ software. Importantly, each cell line displayed a characteristic color depending on the combination and abundance of diverse glycans. In addition, differences in the shades of green observed in NBT-L2b cells indicated the presence of two populations (Figure 1, lower center panel). I named this novel staining method Glycan Painting.

**Figure 1.**
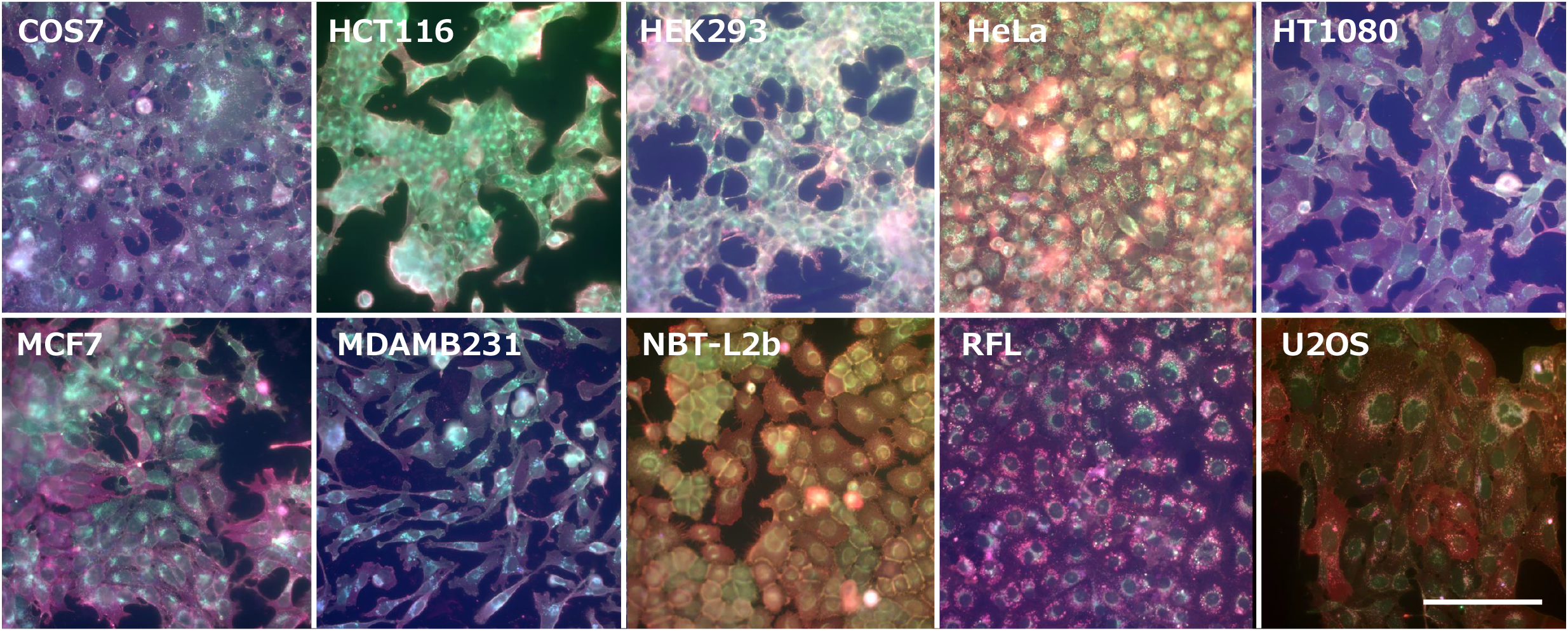
Several representative cell lines stained with Glycan Painting. Each cell was fixed with formaldehyde and ethanol on the coverslips. Cells were stained with ConA-CF405 (blue), WGA-CFR488 (green) and LEL-Dylight 594 (red). Images were captured by a conventional fluorescent microscope equipped with a x40 objective lens and appropriate filter sets, Olympus IX81. All images were adjusted for brightness and contrast using FIJI (Image J) software. Bar: 10 μm.

Next, a mouse embryo section (E14.5) was stained using the Glycan Painting method. Images stained with each fluorescent lectin are shown in Supplementary Figure 2. Glycans are present in all cells of the mouse embryonic body; therefore, all cells can be stained with lectins without exception. In other words, as demonstrated by fluorescent lectin staining images, the ubiquitous presence of glycans across tissues enables lectins to provide broad cellular labeling, in contrast to a marker-dependent approach based on antibodies that exclude unlabeled populations. The intensity and pattern of staining varied across mouse embryonic organs stained with fluorescent lectins (Supplementary Figure 2), indicating that glycan profiles are organ-specific, as previously reported (Otaki et al. 2022). Although a single fluorescent image can only show the distribution of a specific glycan among organs, merging three images provides more information. Figure 2 shows full-color images of mouse embryo sections, demonstrating that different organs and tissues are represented by distinct colors. Furthermore, high-magnification views of the boxed regions in the overall image indicate that each cell type within the tissue is stained in distinct colors by Glycan Painting (Figure 2, right panels). Higher-magnitude views of the selected regions are shown in Supplementary Figure 3. The brain and nerve systems are indicated by dark blue-green with yellow-brown and red dots. Panel 01 shows a high-magnitude image of the midbrain roof, in which the surface region consists of an unformed skull and skin, and cells in various tissues exhibit distinct colors from the surface to the interior. Furthermore, a population of yellow cells was observed among nerve cells in the same regions, although their identity remains unknown (Supplementary Figure 3, panel 01, white arrows). Panel 03 shows a choroid plexus, which is known to consist of choroid epithelium and capillaries. Cells on the surface of the embryonic choroid plexus stained with Glycan Painting appears yellow, with outlines composed of pink and dark blue. The internal region of the choroid plexus contains dark green and blue cells, as well as a small population of red cells (Supplementary Figure 3, panel 03). Panels 20, 22, 24, and 26 show developing bone structures in the mouse embryo. These panels exhibit a light-green mesh-like structure with scattered light-blue punctate signals (Supplementary Figure 3, panel 20). Panel 21 shows the thymus gland, including circular cells delineated with red, blue, and green borders, while a higher-magnitude image allows for better visualization (Supplementary Figure 3, panel 21). Panels 36 and 38 show bilateral sections of the superior mesenteric artery, including red circular cells. Panels 28 and 37 are the midgut, and panel 41 is the gastro-duodenal junction, which exhibits a hierarchical structure with distinctive colors. Panel 35 in Figure 3 and Supplementary Figure 3 show the embryonic lung, including blue immature alveolar tissues and red line structures enclosed by surrounding green outlines. Panels 29 and 30 show liver tissues comprising hepatocytes and non-parenchymal cells. Hepatocytes appear in red, orange, and yellow, indicating different combinations of glycans within this cell type, whereas irregularly shaped blue cells surround them (Supplementary Figure 3, panel 30, white arrows). The cytoplasm was stained red, and cells are likely Kupffer cells (dashed circles). In addition, the upper side of the liver tissue labeled the green region with a yellow line is the diaphragm. Panel 39 in Figure 3 and Supplementary Figure 3 show the pancreatic primordium, with blue or red cells arranged in a honeycomb-like pattern surrounding unclear green outlines. Although Glycan Painting does not directly assign molecular identities to specific cell types, it clearly distinguishes distinct cell populations through differential color signatures.

**Figure 2.**
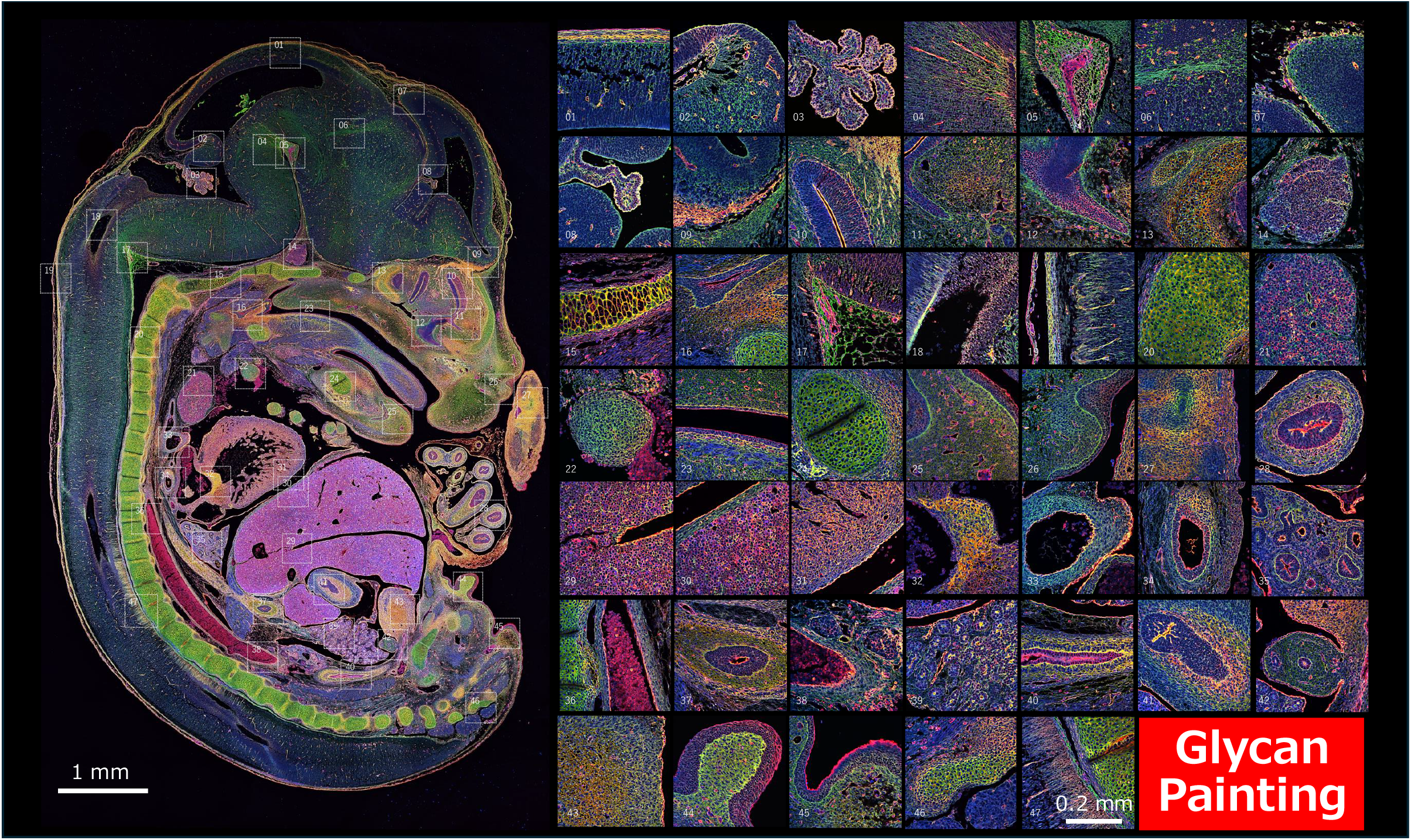
Glycan Painting images of a mouse embryo tissue section. A tissue slice of a mouse embryo (E14.5) was stained with ConA-CF405 (blue), WGA-CFR488 (green) and LEL-Dylight 594 (red). A total of 104 images from a mouse embryo (E14.5) were acquired by a Nikon confocal microscope AX with a x20 objective lens, and individual fluorescence images were stitched into a single composite image by NIS-Elements software. High-magnification images were acquired with a x60 objective lens. Image contrast and color balance were adjusted by FIJI software.

**Figure 3.**
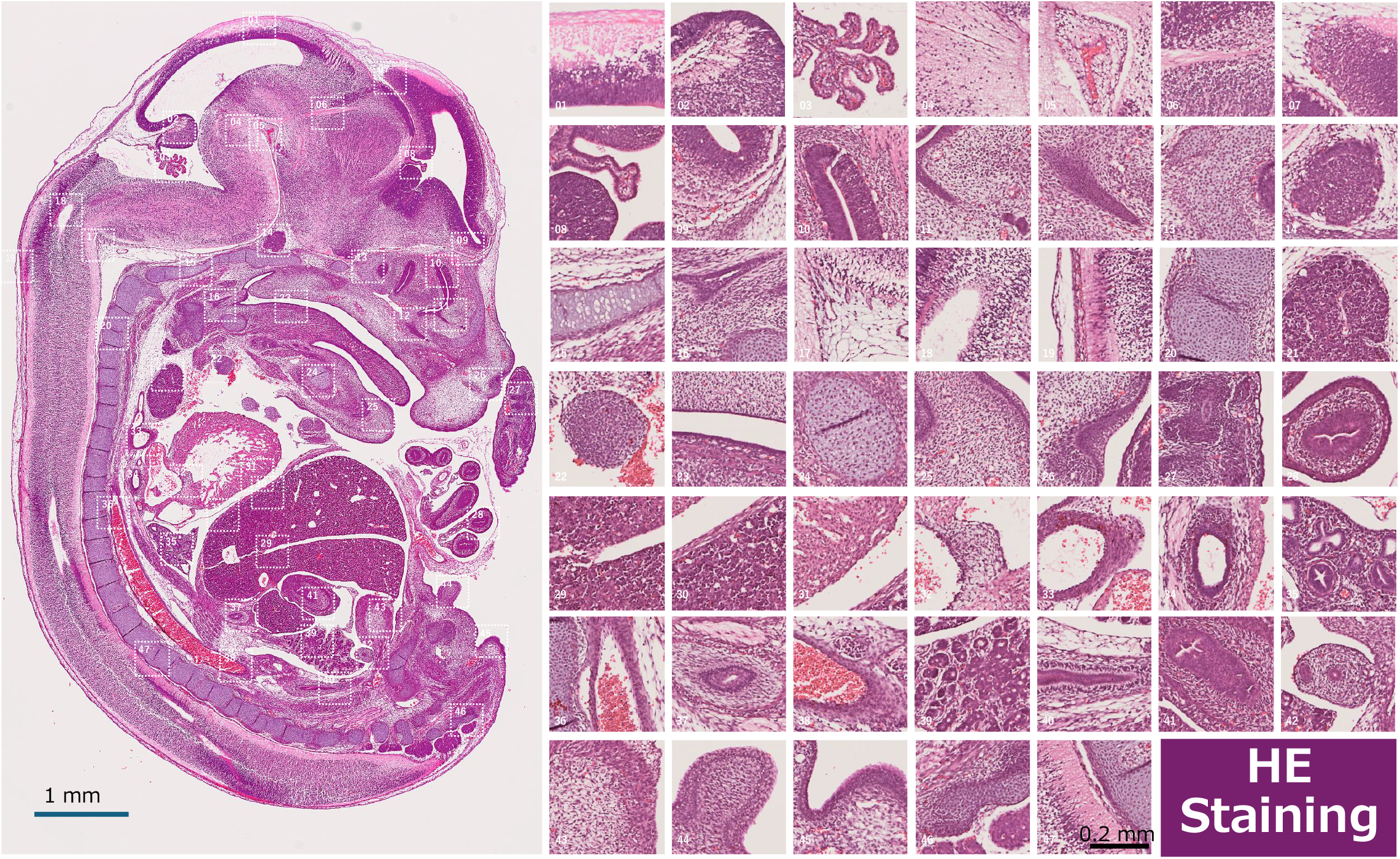
H&E staining of a mouse embryo tissue section. A tissue section of a mouse embryo (E14.5) was stained with hematoxylin and eosin (H&E) solution. H&E images were acquired using a NanoZoomer slide scanner S60 (Hamamatsu Photonics) with a x20 objective lens. High-magnification images were cropped from the original images and enlarged using FIJI software.

Figure 3 shows an H&E-stained serial section adjacent to the Glycan Painting section from the same paraffin-embedded block. H&E staining is a gold standard among histological methods in which hematoxylin and eosin stain nuclei blue-purple and cytoplasm pink-red, respectively, enabling the assessment of tissue morphology. Similar to the H&E staining images (Figure 3), Glycan Painting also allows a general but comprehensive evaluation of the overall morphology of tissues. In addition, because glycans exist in all cells, this method visualizes all cell populations through staining with lectins targeting ubiquitously expressed glycans. While H&E staining provides morphological information, Glycan Painting integrates a morphological overview with color-based cellular discrimination in a single image.

A previously developed and related method, Actin Painting (Nagasaki et al. 2025), also allows several cell types in a tissue to be visualized as distinct colors. Actin Painting is a triple-fluorescence staining with three types of actin probes, such as phalloidin and Lifeact, conjugated to fluorescent dyes. However, most actin probes fail to bind denatured actin filaments and are therefore not applicable to formalin-fixed, paraffin-embedded sections. In contrast, because Glycan Painting targets glycans within cells, it can be applied to strongly fixed samples and is not affected by antigen inactivation, making it a highly versatile technique. To date, substantial efforts worldwide have focused on identifying specific markers and developing corresponding probes for target cell identification. To address this need, Glycan Painting is based on triple staining with typical fluorescent lectins and is a simple and broadly accessible technique. Furthermore, unlike cyclic immunofluorescence methods, which rely on iterative staining and stripping cycles to achieve high multiplex imaging, Glycan Painting accomplishes full-color imaging in a single staining step.

Finally, the micrographs presented in this report demonstrate the comprehensive labeling of cells within tissue sections based on cell-type-specific combinations of glycans. This approach enables full-color imaging of all cells within a living organism, which was not previously achieved in multiplex fluorescence imaging. The assignment of distinct colors to individual cell types provides an intuitive framework for cell identification. Furthermore, Glycan Painting distinguishes cell populations by assigning cell types as a unique color signature, enabling the spatial mapping of cell types defined in cell atlases onto tissue sections, thereby bridging transcriptome-based cell classification with histological architecture in the future. If specific color signatures can be linked to defined cell types, cell identities can be directly represented by color-based information. Importantly, a major advantage of this method is its use of readily available reagents and is compatible with standard fluorescence microscopes, making it broadly accessible.

## Abbreviations

ConA: concanavalin A
H&E: hematoxylin and eosin
LEL: *Lycopersicon esculentum* (tomato) lectin
PBS: phosphate-buffered saline
WGA: wheat germ agglutinin
TBS: Tris-buffered saline.

## Acknowledgements

This research was supported by Management Expenses Grants from AIST, AMED (Grant Number JP23ym0126803), and the Japan Society for the Promotion of Science (JSPS) under KAKENHI (Grants) 24K22414 and 25H01226. The author is grateful to K. Hirano and C. Okatani for helpful feedback on this paper.

---

**Supplementary Figure 1. Conceptual Schematic Overview of Glycan Painting**.

(A)Schematic illustration of the principle of Glycan Painting.

The combination of glycan species within cells is cell-type-specific, and three lectins labeled with different fluorescent dyes (red, green, blue) bind to cells according to their glycan composition. By merging the fluorescence images from each fluorescent lectin, a color image can be generated, in which colors of each cell type are determined by the RGB ratios of the individual fluorescent lectins.

(B)Additive color wheel diagram. This diagram indicates that mixing two of the three primary colors (red, green, and blue) generates the secondary colors, cyan, magenta, and yellow, whereas mixing all three prime colors produce white. In practice, fluorescent lectins bind to cells in various combinations, producing full-color composite images.

**Supplementary Figure 2. Monochromatic images of a tissue slice from a mouse embryo stained with Lectin Painting**.

The tissue slice was stained simultaneously with ConA-CF405 (blue), WGA-CFR488 (green) and LEL-Dylight 594 (red). Each image was acquired using the Nikon AX microscope. Merged images of them were indicated in Figure 3. Bars; 1 mm and 0.2 mm.

**Supplementary Figure 3. Higher-magnified view of a mouse embryo section stained using Glycan Painting**.

Higher-magnitude views of selected regions from Figure 2. midbrain roof of a mouse embryo (Panel 01), choroid plexus (Panel 03), developing bone (Panel 20), thymus gland (Panel 21), midgut (Panel 28), liver (Panel 30), lung (Panel 35), superior mesenteric artery (Panel 36), and pancreatic primordium (Panel 39).

## Notes

### Competing Interest Statement

The authors have declared no competing interest.

