## Supplemental Files for "Glycan Painting: Triplex Lectin Staining Enables Visualization of Cell-Type-Specific Glycan Profiles in Tissue Sections"

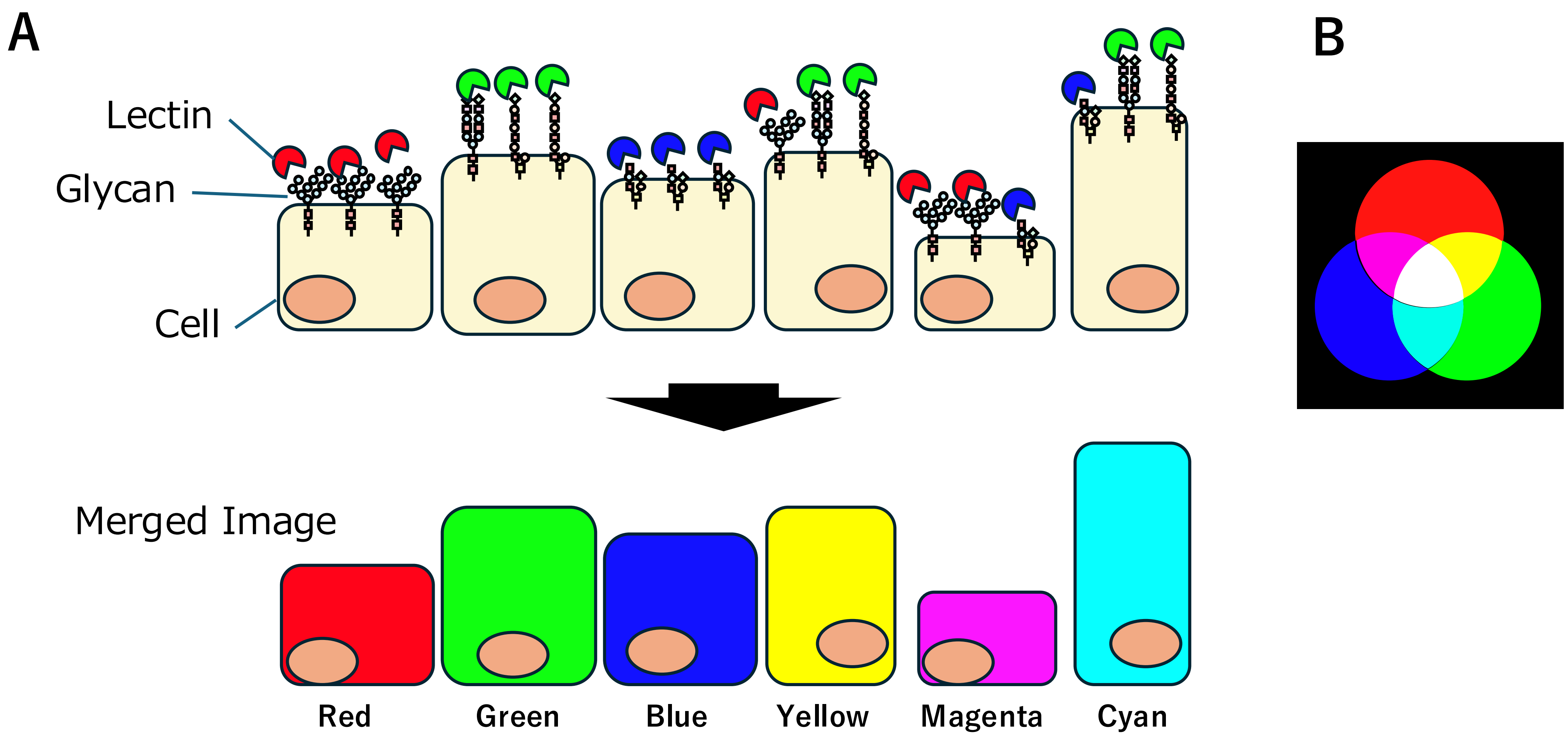


Nagasaki Supplementary Figure 1

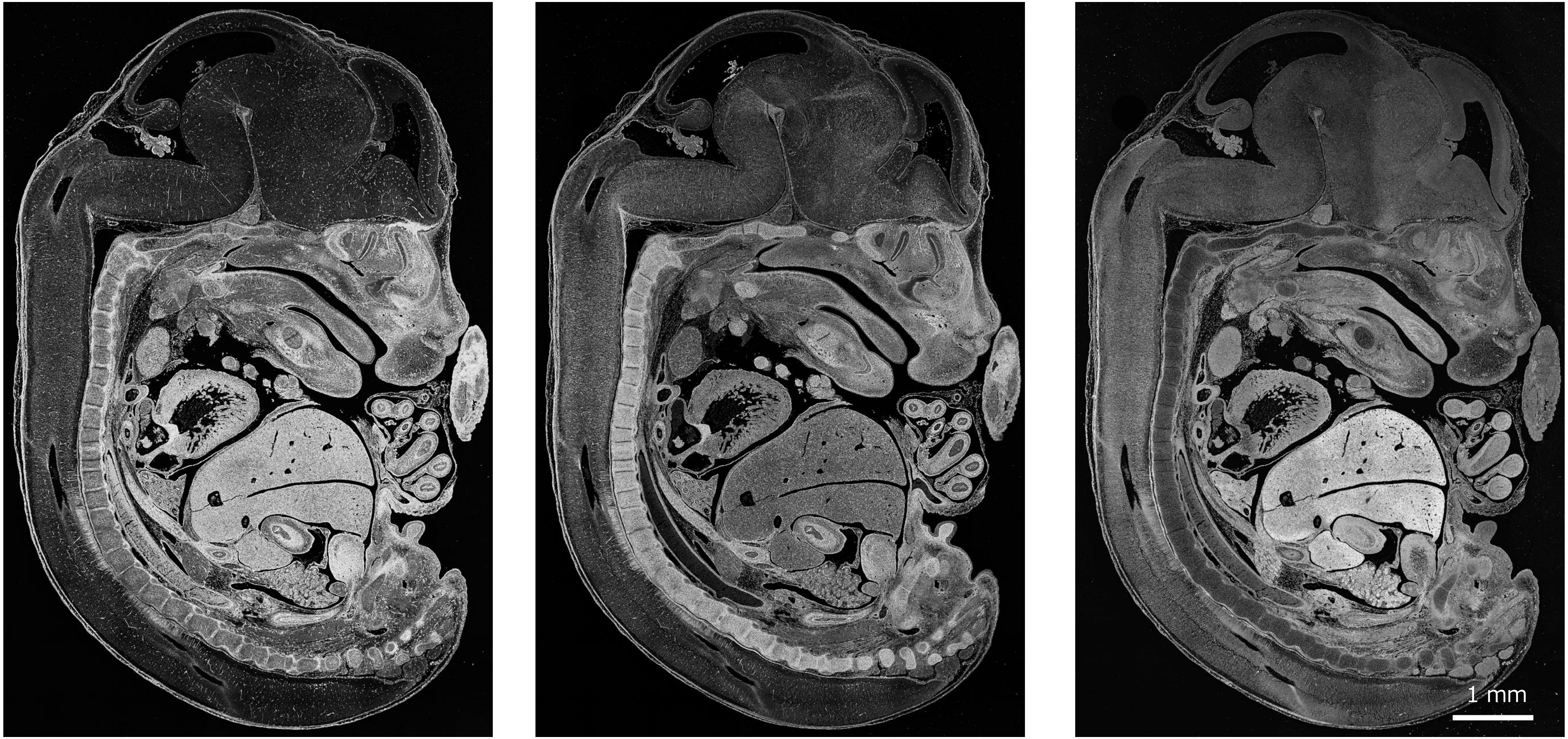


TL-Dylight 594 WGA-CFR488 ConA-CF405

Nagasaki Supplementary Figure 2

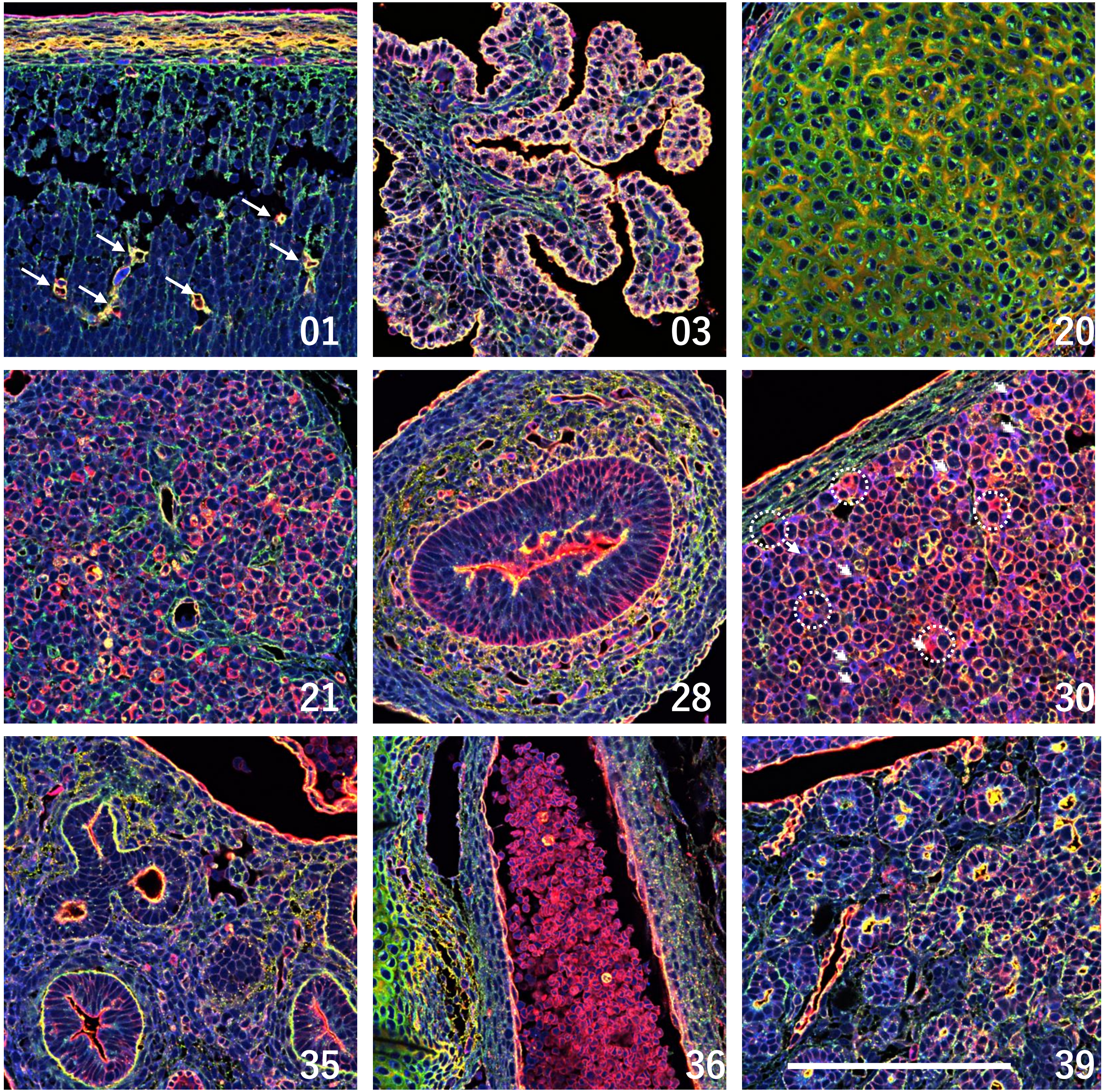


Nagasaki Supplementary Figure 3
